# Investigation of Double-Stranded DNA Donors and CRISPR-Cas9 RNP for Universal Correction of Mutations Causing Cystic Fibrosis in Human Airway Cells

**DOI:** 10.1101/2025.09.21.677613

**Authors:** Vrishti Sinha, Paul G. Ayoub, Colin J. Juett, Lindsay E. Lathrop, Ruth A. Foley, Ruby A. Sims, Joseph D. Long, Emily C. Duggan, Neil Fernandes, Beate Illek, Brigitte N. Gomperts, Steven J. Jonas, Donald B. Kohn

**Affiliations:** Department of Human Genetics, University of California, Los Angeles; Los Angeles, CA 90095, USA; Molecular Biology Institute, University of California, Los Angeles; Los Angeles, CA 90095, USA; Department of Microbiology, Immunology, and Molecular Genetics, University of California, Los Angeles; Los Angeles, CA 90095, USA; Department of Bioengineering, University of California, Los Angeles, Los Angeles, California 90095, United States; Department of Pediatrics, David Geffen School of Medicine, University of California, Los Angeles, Los Angeles, California 90095, United States; Department of Pediatrics, School of Medicine, University of California, San Diego, San Diego, California 92103, United States; Jonsson Comprehensive Cancer Center, University of California, Los Angeles, Los Angeles, California 90095, United States; Eli & Edythe Broad Center of Regenerative Medicine and Stem Cell Research, University of California, Los Angeles, Los Angeles, California 90095, United States; California NanoSystems Institute, University of California, Los Angeles, Los Angeles, California 90095, United States

**Keywords:** Cystic Fibrosis, Site-specific Insertion, Nonviral gene delivery, Homology-Directed Repair, Cystic Fibrosis Transmembrane Conductance Regulator (CFTR)

## Abstract

Cystic fibrosis (CF) is a devastating genetic disease caused by mutations in the *Cystic Fibrosis Transmembrane Conductance Regulator* (*CFTR*) gene. As morbidity and mortality from CF results from a lack of mucus clearance that leads to chronic bacterial infections and progressive loss of lung function, site-specific insertion of a *CFTR* cDNA into the endogenous *CFTR* locus in airway basal stem cells (ABSCs) could prove curative for all disease-causing mutations. This study describes the development of nonviral genome editing reagents, designed to be packaged into nonviral delivery systems. An sgRNA targeting the 5’ untranslated region of *CFTR* was characterized as directing high on-target cutting and displaying a safe off-target profile. Airway cell lines electroporated with chemically-modified (1-Aminohexane - AmC6), linear double-stranded DNA (ldsDNA) constructs were utilized as an Homology Directed Repair (HDR) donor, initially optimized with an mCitrine reporter. Expectedly, when the 780bp mCitrine cDNA was replaced with the 4.4kb *CFTR* cDNA, integration efficiency dropped significantly. However, 1-2% integration of codon optimized donors was sufficient to restore CFTR expression in the bulk edited population of human bronchial epithelial cell line, 16HBE14o- (16HBE), to levels reaching 50% of wildtype expression as measured by Western Blot. Electrophysiological validation of CFTR ion channel function measured via Ussing Chamber Assay revealed that these bulk edited populations exhibit greater than 40% restoration of the chloride ion currents of the measured wildtype controls. These results demonstrate that low levels of *CFTR* integration can be made therapeutically relevant by optimizing the designs of gene editing reagents. Importantly, this work utilizes nonviral editing reagents, an essential step towards *in vivo* gene therapy for CF.

## Introduction

Cystic fibrosis (CF) is a rare autosomal recessive monogenic disorder resulting from mutations in the *Cystic Fibrosis Transmembrane Conductance Regulator* (*CFTR*) gene which encodes for an epithelial ion channel protein that transports chloride and bicarbonate across the membrane^1, 2^. This transmembrane protein is essential for maintaining proper homeostatic osmosis in the epithelial layers of the lungs, pancreas, bile ducts, intestines, and sweat glands^2^. When mutated, dysfunctional CFTR protein fails to transport chloride across these various epithelia properly such that the mucus layer dehydrates, leading to frequent pulmonary infections and inflammation that ultimately results in progressive lung destruction and respiratory failure^3^. Despite affecting multiple organs, patient morbidity and mortality result primarily from complications in the airway, thereby highlighting the importance of targeting the lungs for corrective gene therapy approaches ^4^.

The use of both corrector and potentiator modulators to improve either the folding or function of the mutated CFTR protein, respectively, has greatly improved the health of many CF patients^5-7^. However, despite the success of modulator therapies, not all patients are able to benefit from these medications, particularly those with null mutations or very low levels of CFTR expression and those who develop side effects from these treatments. Roughly 10% of patients have mutations within these categories, highlighting the need for alternative therapies to maximize the benefits for the entire CF patient population^5, 8^.

The genome editing capabilities enabled by the CRISPR/Cas9 system have allowed for site-specific repair or replacement of mutated genes in a wide variety of disease contexts. Consequently, for *CFTR*, stable integration of a normal *CFTR* cDNA into the endogenous *CFTR* locus in airway basal stem cells (ABSCs) downstream of the promoter has proven beneficial to restoring expression of functional CFTR levels^9^. Since this approach is mutation-agnostic, it represents a universal approach that can be functionally curative for all CF patients, regardless of the mutation type.

The limiting factor of this approach is the large size of the coding region of the *CFTR* gene. At roughly 4.4kb, traditional homology directed repair (HDR) donors carrying the *CFTR* coding sequence exceed the packaging capacity of a single adeno-associated virus type 6 (AAV6). As a result, traditional AAV-based HDR approaches have shown low efficiency in delivering the necessary coding sequence^10, 11^. Some groups have also targeted high frequency mutations with base and prime editing, achieving partial functional correction of the CFTR protein^12-14^. Still, with over 2,000 known *CFTR* mutations, these strategies will leave many patients without an option for a curative gene therapeutic approach^15^. Novel strategies to address the size limitation have emerged by generating shortened *CFTR* variants that package into a single AAV6 vector^16^ or by delivering the full-length coding sequence of *CFTR* in segments using AAV6 vectors. In the latter approach, edited airway basal cells were enriched *ex vivo* and retained their phenotype, restoring CFTR function near wildtype cells as measured *via* Ussing Chamber Assay^17, 18^.

Still, the necessity of both enrichment of edited cells and the use of AAV6 as an HDR donor requires the editing to be performed *ex vivo* (**Figure 1A**). Apart from editing efficiencies concerning large donors, the process of autologous transplant for airway basal cells is limited by complex roadblocks which have not yet been overcome^19, 20^. Alternatively, *in vivo* approaches could rely on the nonviral delivery of genome editing reagents directly to the patient’s lung using nanoparticle-based carriers, thereby potentially overcoming the limitations of autologous transplantation (**Figure 1A**). Furthermore, this *in vivo* editing approach would be less invasive to the patient and avoid the potential innate immune response that may occur in response to viral vectors.

**Figure 1.**
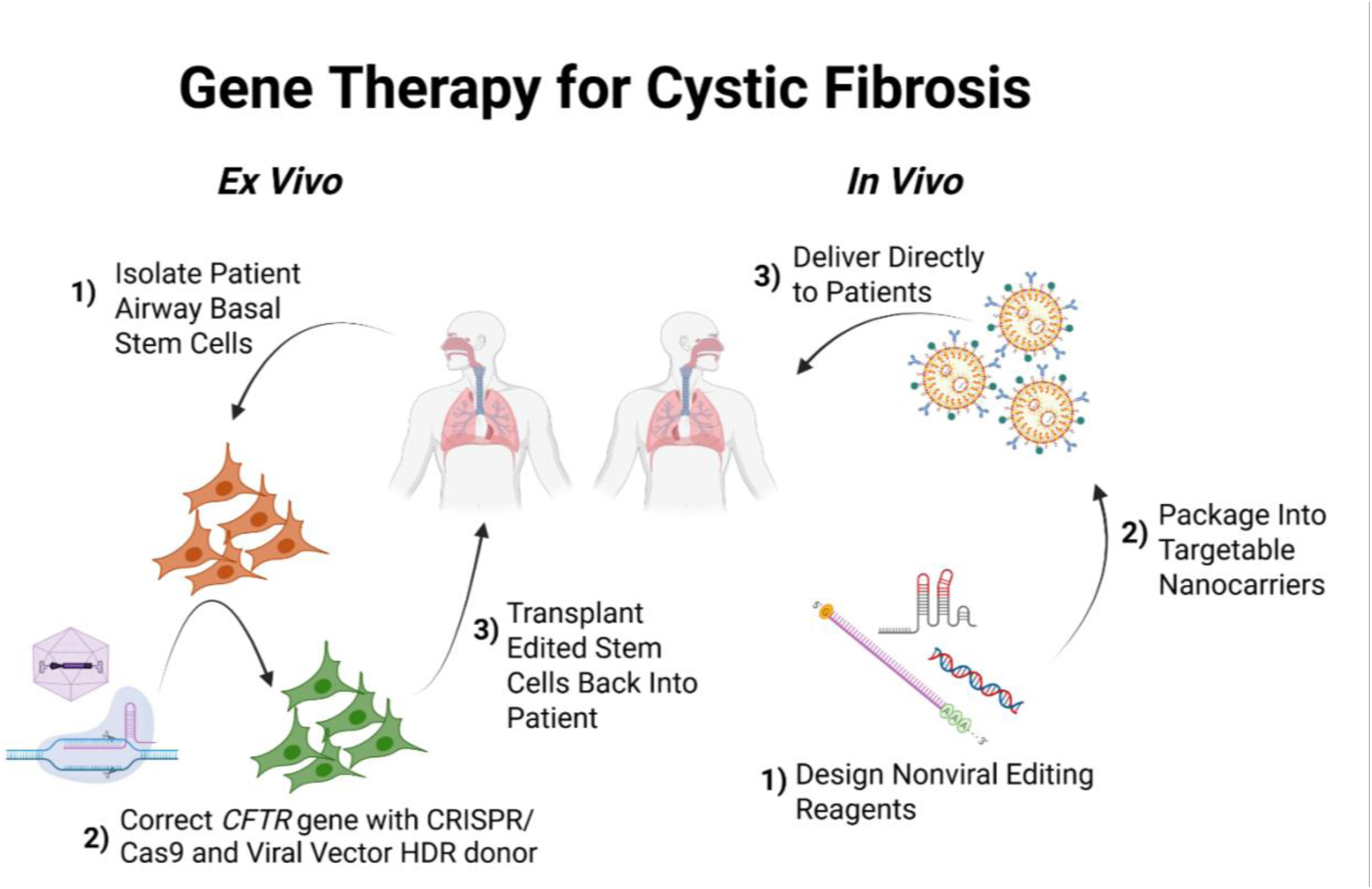
Gene Therapy for Cystic Fibrosis. Description of two gene therapy approaches for Cystic Fibrosis achieved via *ex vivo* or *in vivo* genome editing. Created with Biorender.

Due to these benefits, we sought to develop a nonviral genome editing system to insert the *CFTR* cDNA downstream of the endogenous *CFTR* promoter that could be optimized for *in vivo* delivery. We aimed to override the majority of coding region mutations and acquire appropriate expression of CFTR to reduce disease phenotype, which is expected to be achievable with about 10% correction^21^. Here, we report an editing system that utilized CRISPR/Cas9 to insert a codon-optimized *CFTR* cDNA cassette into the 5’ Untranslated Region (UTR) of the *CFTR* gene. Using this system, CFTR protein was successfully produced in a CFTR-null bronchial epithelial cell line even at low (1-2%) integration efficiencies. When edited cell populations were grown as submerged cell monolayers in Transwell membrane inserts, and assessed for functional restoration, chloride currents of up to 80% that of wildtype were achieved. These results demonstrate a step towards a mutation agnostic nonviral gene editing approach to stably treat CF.

## Results

We began our studies by investigating the optimal site in the endogenous *CFTR* locus to insert a *CFTR* cDNA cassette. As such, various single guide RNAs (sgRNA) targeting either the 5’UTR or Intron 1 of the *CFTR* gene were screened for frequency of target site disruption (**Figure 2A**). Initially, guide screening was performed in a T84 cell line derived from a human lung metastasis of colon carcinoma. Screening was repeated in the 16HBE cell line, which is more conducive for downstream functional assays and therefore a more clinically relevant model than T84 cells (**Figure 2B**). All guides (**Supplemental Table 1)** were screened with the addition of the Alt-R Cas9 Electroporation Enhancer to assist in increasing *ex vivo* editing efficiency, thereby achieving a two-fold increase in CRISPR-Cas9 cutting efficiency (**Supplemental Figure 1A, 1B**). In edited T84 cells, sgRNA 3 targeting the 5’UTR and delivered as sgRNA-Cas9 ribonucleoprotein (RNP) had an average cutting frequency of 35.7% as measured by the formation of insertion-deletions (indels) (**Figure 2C**). In comparison, the Intron 1 guides (sgRNA13-19) had average cutting efficiencies ranging from 65% up to 93.3% (**Figure 2C**).

**Figure 2.**
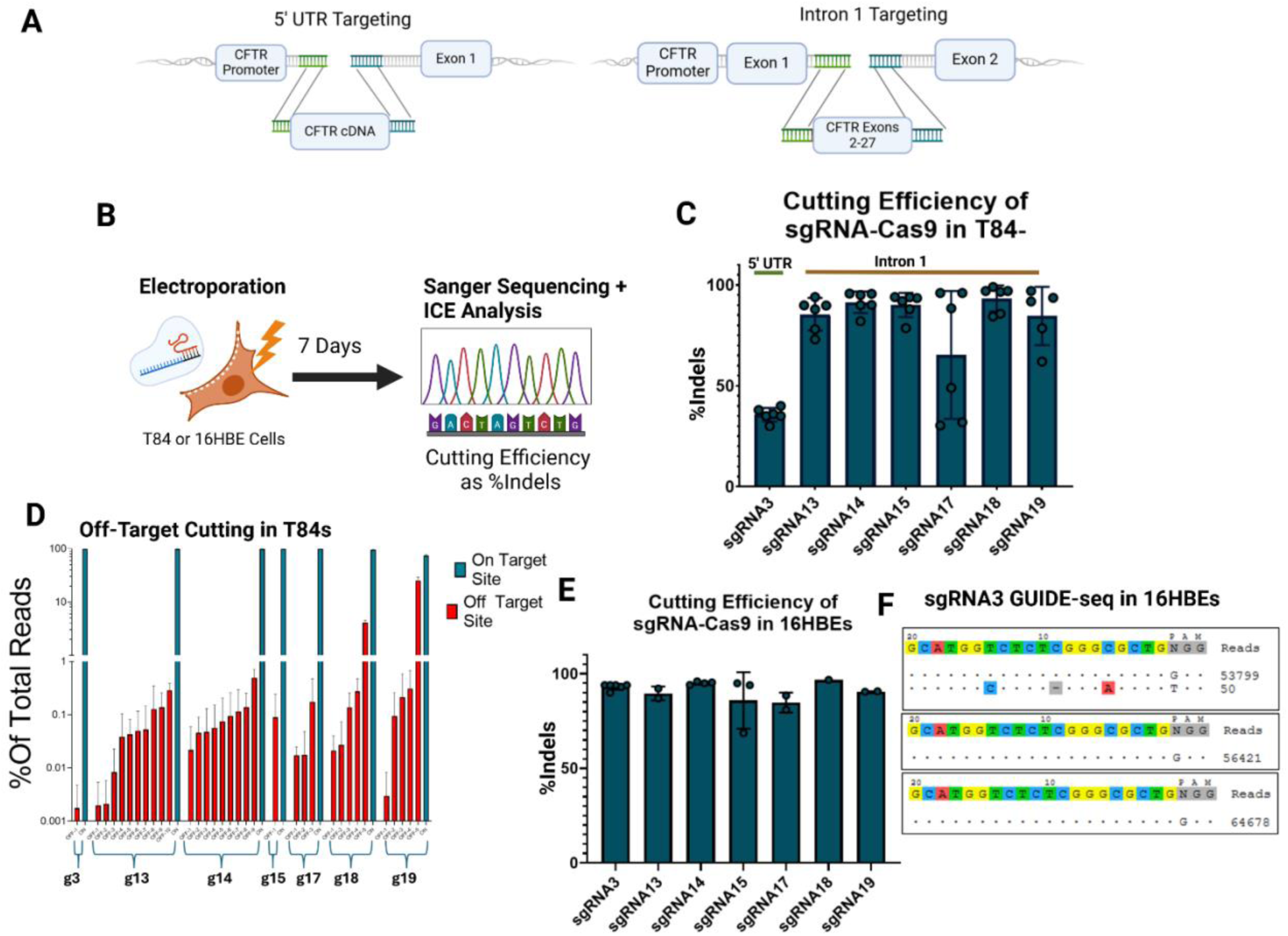
Characterizing sgRNA Targeting *CFTR* Locus. (A) Schematic demonstration of two editing strategies based on two sgRNA targets. (B) Method timeline for screening sgRNA-mediated cutting efficiency. (C) Cutting efficiency, measured by % indels, in T84 cells (n=6). Each dot represents a different electroporated replicate. (D) Percent of total reads that are on-target (blue) or off-target (red) from the highest dose of dsODN delivered (5µM) in GUIDE-seq analysis (n=3). Different red bars represent distinct sites. (E) Cutting efficiency, measured by %indels, in T84 cells (n=2-6). Each dot represents a different electroporated replicate. (F) Results from GUIDE-seq analysis of sgRNA3 in 16HBEs from the highest dose of dsODN delivered (5µM) in GUIDE-seq analysis (n=3). Statistical significance was analyzed using a one-way ANOVA followed by multiple paired comparisons for normally distributed data (Tukey test). All statistical tests were two-tailed and a p value of < 0.05, **p < 0.01, ***p < 0.001, ****p < 0.0001. (A,B) created with BioRender.

The off-target activity of sgRNA-directed Cas9 was assessed *via* GUIDE-seq in the T84 cell line^22,23^. Although off-target reads were observed from all screened guides, the 5’UTR sgRNA 3 and the Intron 1 sgRNA 15 had the fewest measured off-target cut sites, and these were rare with all comprising less than 0.1% of total reads (**Figure 2D**).

In follow-up guide screening studies in the more clinically relevant 16HBE cells, the cutting efficiency of sgRNA 3 dramatically increased to 93.2% compared to its lower activity seen in the T84s, while the Intron 1 guide maintained high cutting efficiency in both cell lines, with up to 96.8% indels in the 16HBE cells (**Figure 2E**). This result highlights the differences in guide efficiency across cell lines.

The off-target activity of sgRNA 3 in 16HBE cells was analyzed via GUIDE-seq, which revealed a single off-target site in an intron of *POLR2J2* (**Figure 2F**). This gene encodes for a subunit of DNA-Dependent RNA Polymerase II, which is not involved in cell cycle regulation, and therefore is unlikely to lead to oncogenesis. Due to the increased safety profile of sgRNA 3 and its potential to treat mutations found within Exon 1 (which would not be addressed with the downstream Intron 1 sgRNA), sgRNA 3 was selected to test HDR donor reagents since the entire *CFTR* cDNA could be inserted into the 5’UTR.

Next, nonviral DNA constructs were tested as HDR donors. To first assess donor integration potential, an mCitrine construct was designed and tested to target the 5’UTR cut site of sgRNA 3 with flanking Homology Arms (HA) of roughly 500bps. To improve expression, bovine growth hormone polyA signal (BGA) sequence was included in the reporter construct as well. In 16HBEs, this construct was delivered as either plasmid forms or as ldsDNA HDR donors. The safety and efficacy of these donors were assessed by delivery combined with sgRNA3-Cas9 RNP, followed by an 3-(4,5-dimethylthiazol-2-yl)-2,5-diphenyltetrazolium bromide (MTT) Assay^24^ to determine relative cell viability, and an in-out digital droplet PCR (ddPCR) assay on genomic DNA of edited cells to determine allelic integration frequencies (**Figure 3A, 3B**).

**Figure 3.**
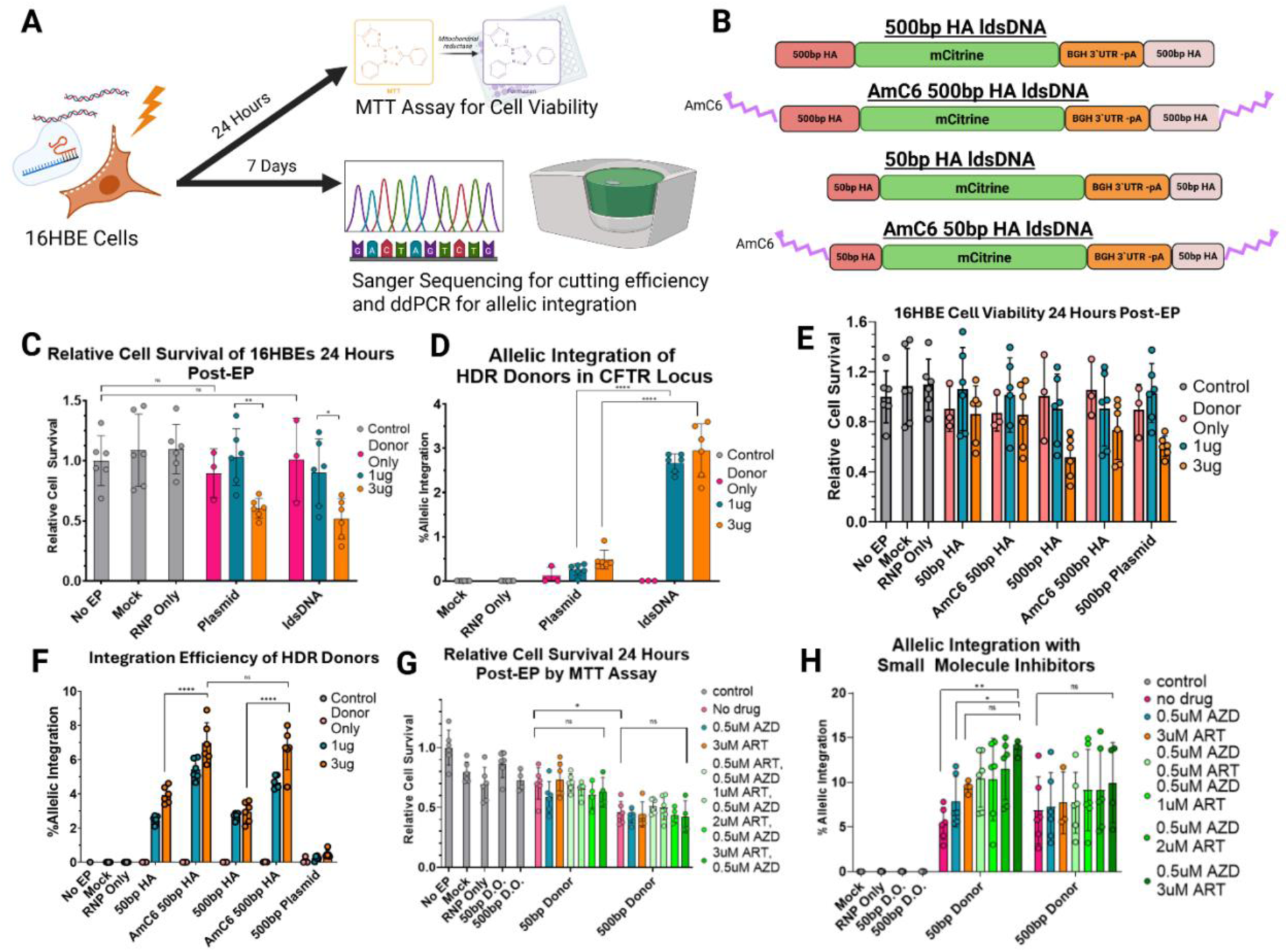
Optimization of Nonviral HDR Donor Editing Reagents. (A) Illustration of workflow for screening sgRNA-mediated cutting efficiency. Cells were divided immediately following electroporation to perform MTT assay and molecular analysis separately (B) Schematics for mCitrine HDR donors. All donors contain mCitrine gene and 3’UTR if the bovine growth hormone gene. Donors varied based on homology arm size (50bp or 500bp) and inclusion or exclusion of AmC6 modification. (C) Relative cell survival and (D) integration after electroporation delivering plasmid or linear dsDNA HDR donor with 500bp HA construct to 16HBE cells (n=3). (E) Relative cell survival and (F) integration after electroporation delivering 50bp homology arm donors with or without AmC6 and 500bp donors with or without AmC6 (n=3). (G) Relative cell survival and (H) integration after electroporation delivering AmC6 50bp HA and AmC6 500bp HA donors with varied concentrations of AZD7648 and ART558 in recovery media (n=3). Each dot represents a different electroporated replicate. (F) Results from GUIDE-seq analysis of sgRNA3 in 16HBEs from the highest dose of dsODN delivered (5µM) in GUIDE-seq analysis (n=3). Data are represented as mean ± SD of biological triplicates from three experiments. Statistical significance was analyzed using a one-way ANOVA followed by multiple paired comparisons for normally distributed data (Tukey test). All statistical tests were two-tailed and a p value of < 0.05, **p < 0.01, ***p < 0.001, ****p < 0.0001. (A,B) created with BioRender.

While delivery of the donors on their own did not significantly affect cell viability, significant toxicity was observed in the presence of RNP in combination with the highest doses of both ldsDNA and plasmid HDR donors (**Figure 3C**). Despite similar toxicity profiles, the ldsDNA dramatically outperformed plasmid in allelic integration frequency, with over six-fold greater integration (**Figure 3D**).

Nanoplasmids were also tested for HDR donor delivery in T84 cells and demonstrated significantly less toxicity than the plasmid HDR donor (**Supplemental Figure 2A**). However, the nanoplasmid exhibited significantly lower integration efficiencies compared to ldsDNA, with no significant difference compared to the plasmid HDR donor (**Supplemental Figure 2B**).

It is well-established that HDR donor size inversely correlates with integration efficiency^25, 26^. The ldsDNA donor encoding for the mCitrine cassette, which is roughly 2.0kb, achieved 3.0% integration. In contrast, the *CFTR* cDNA alone, without necessary homology arms or regulatory elements, is roughly 4.4kb. Given our aim to reach the therapeutic threshold of 6-10% targeted integration, optimization of modifications and reagents to boost transgene integration was necessary prior to testing donors containing the *CFTR* cDNA.

To enhance HDR efficiency, the initial ldsDNA donor were designed with larger 500bp homology arms that included the *CFTR* promoter sequence on the 5’ end. However, inclusion of the promoter sequence raised concerns about potential dysregulation of nearby genes due to off-target integration events or random, nonspecific trapping of the donor. To mitigate this risk, we designed and tested HDR donors with shorter 50bp homology arms that excluded the *CFTR* promoter sequence and evaluated whether efficient integration could still be achieved.

Furthermore, chemical modifications on the 5’ ends of ldsDNA donors, such as 1-Aminohexane (AmC6), have been shown to boost gene knock-in rates up to fivefold, and permit the use of shorter homology arm lengths with minimal loss of integration efficiency (**Figure 3B**)^27^. We sought to assess the effect AmC6 modifications on the 5’ ends of the donor construct in combination with the reduced homology arm length.

In edited 16HBE cells, the relative cell survival was unchanged with the addition of the AmC6 modification (**Figure 3E**). Encouragingly, the allelic integration increased approximately twofold with the AmC6 modification across both 50bp and 500bp donor homology arms (**Figure 3F**). Furthermore, reduction in the size of the homology arms did not significantly change the integration rate, supporting the use of the 50bp homology arm HDR donor (**Figure 3E**). Notably, in the T84 cell line, the addition of AmC6 to the donors had a modest increase in integration. Moreover, reducing homology arm length in this context decreased knock-in efficiency (**Supplemental Figures 2C, 2D**). These findings suggest the benefit of the AmC6 modification might be cell type specific.

To support efforts to increase site-specific integration of transgenes, there have been several recent studies into small molecule inhibitors acting on the DNA-dependent protein kinase (DNA-PK) catalytic subunit (AZD7648 - AZD) and Polymerase Theta (ART558 - ART), two important mediators of the nonhomologous end-joining (NHEJ) and microhomology mediated end-joining (MMEJ) pathways, respectively^28, 29^. To clarify whether the 500bp donor might have an improved propensity for HDR, both the 50bp and 500bp HA donors with the AmC6 modification were tested head-to-head in a drug titration. Cell survival was unaffected by the addition of the drugs at any concentration or combination, but the 50bp HA donor supported significantly better cell survival than the 500bp HA donor (**Figure 3G**).

Surprisingly, the 500bp HA donor had no significant differences in integration with the addition of the inhibitors. However, the 50bp HA donor had significantly improved integration, doubling to an average of 13.80% knock in efficiency at the highest combined dose of AZD and ART (**Figure 3H**). Moreover, the combination of the drugs supported significantly higher integration than AZD on its own (**Figure 3H**). From conducting these optimization experiments, the optimal candidate ldsDNA donor included 50bp homology arms, AmC6 modifications on the 5’ ends, and the addition of 0.5uM AZD and 3uM ART into the recovery medium.

With the optimal editing HDR delivery system identified, the mCitrine cDNA was exchanged for the full-length *CFTR* cDNA. This construct included AmC6-modified homology arms of 50 bp, and Woodchuck Posttranscriptional Response Element (WPRE), and Bovine Growth Hormone (BGH) polyadenylation signal sequence. Codon optimization, the process of altering synonymous codons to better align with host-specific expression preferences, can improve translation efficiency, mRNA stability, and protein yield when designing candidate constructs, however, algorithms differ in their approaches, thus potentially affecting downstream expression and function^30-32^. Three codon-optimized *CFTR* constructs were therefore designed using the publicly available Java Codon Adaptation Tool (JCAT)^33^, GeneArt (GA)^34^, and IDT algorithms (**Figure 4A)**. These donors were evaluated and compared in the 16HBEge-G542X cell line^35^. This cell line, created by the Cystic Fibrosis Foundation, has the G542X *CFTR* mutation knocked into the endogenous site. This early stop codon mutation causes transcribed mRNA to be degraded through the nonsense-mediated decay pathway, resulting in a lack of CFTR protein production.

**Figure 4.**
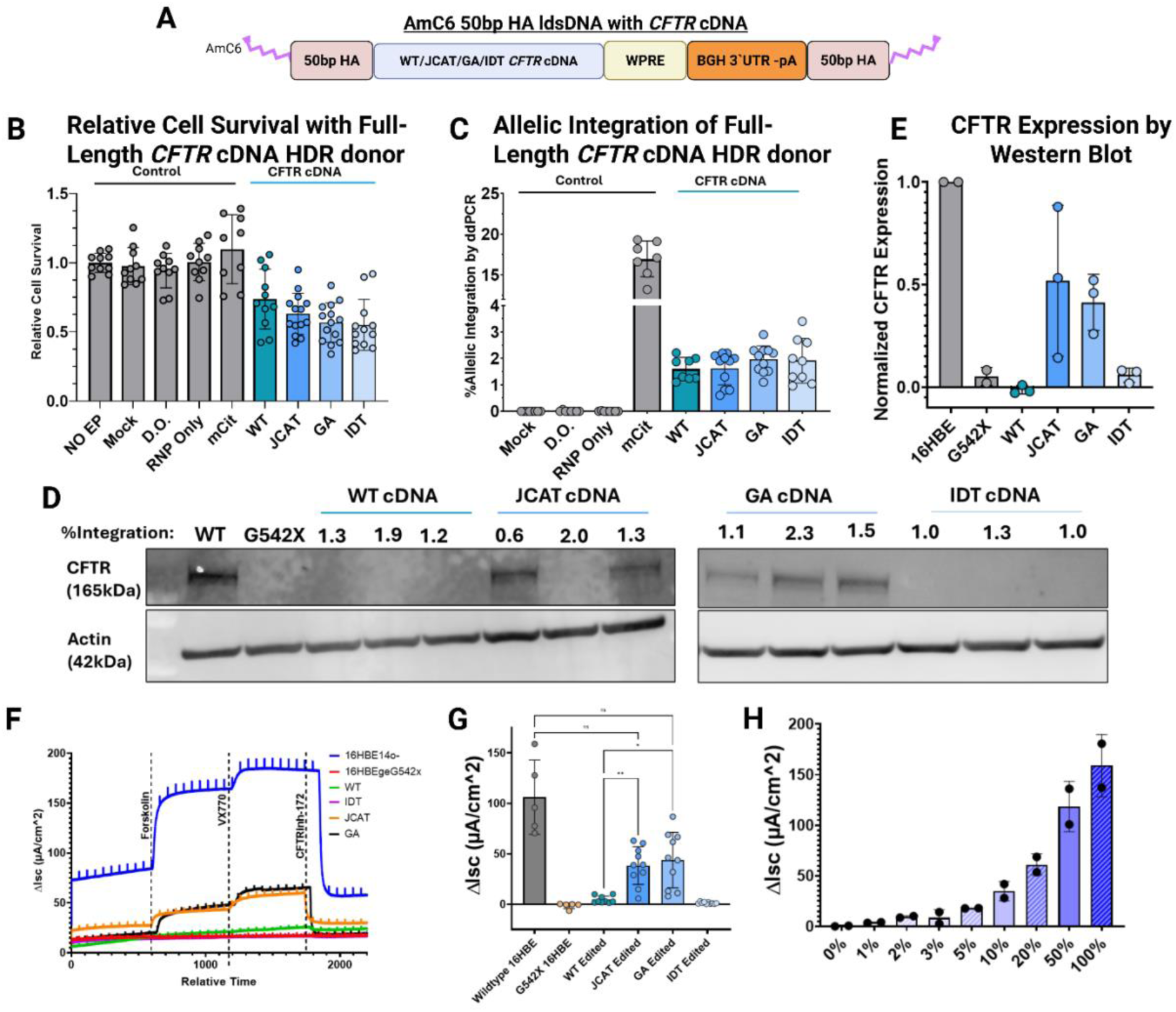
Editing With *CFTR* cDNA Achieves Expression and Function. (A) Schematic of CFTR ldsDNA donors (B) Relative cell survival and (C) integration of 16HBEgeG542X cells after electroporation delivering AmC6 50bp HA HDR donors carrying the wildtype and 3 codon optimized *CFTR* cDNA (n=3). (D) Western Blot from lysate collected from wildtype, G542X, and bulk edited cell population. Percent integration is given below the HDR donor name. (E) CFTR protein levels were quantified by densitometry using ImageJ and normalized to Actin levels (n=3). (F) Representative Ussing Chamber traces. (G, H) Maximum inhibitable transepithelial chloride current values measured in (G) edited 16HBEge-G542X monolayers or (H) mixed populations of wildtype and G542X 16HBE cells at increasing percentages. Statistical significance was assessed using a one-way ANOVA followed by multiple paired comparisons for normally distributed data (Tukey test). All statistical tests were two-tailed and a p value of < 0.05, **p < 0.01, ***p < 0.001, ****p < 0.0001.

Cells edited with the *CFTR* cDNA donors had significantly increased toxicity compared to the mCitrine donor (**Figure 4B**). This higher cytotoxicity was attributed to the increased size of the *CFTR* donors, as it is known that larger physical sizes of DNA constructs are correlated to increased specific toxicity^36^. Integration frequencies of the larger donors significantly dropped as well, compared to the smaller mCitrine donors, with the average integration for each construct ranging between 1-2% (**Figure 4C**).

CFTR protein expression from the bulk edited populations was assessed *via* Western Blot (**Figure 4D)**. Despite the low integration of the *CFTR* constructs, the bulk-edited cell populations with the JCAT and GA codon optimizations achieved roughly 52% and 41% of the CFTR protein expression as in the WT 16HBE control, respectively (**Figure 4E)**. No measurable expression of CFTR protein was seen for the WT or IDT codon optimized *CFTR* cDNA construct, demonstrating the value of codon optimization in increasing translation of the transgene protein.

The bulk edited 16HBEge-G542X cell populations were seeded in Transwell plates and cultured for 7 days prior to transepithelial chloride current assessment using an Ussing Chamber Assay to verify functional recovery of the CFTR protein (**Figure 4F**). The populations edited with the JCAT and GA HDR donors had an average CFTR-inhibitor172-specific current response of -38.4µA/cm^2^ and -43.9µA/cm^2^, respectfully (**Figure 4G**). Both currents were significantly greater than the 16HBEge-G542X control, or the cell populations edited with the WT HDR donor (**Figure 4G**), thereby achieving functional current recovery to 40% of that of WT levels.

Considering the partial functional recovery it was imperative to confirm that the functional activity from 16HBEge-G542X correlates with functional activity from WT 16HBE cells. Therefore, an Ussing Chamber Assay was performed on monolayers of different ratios of WT 16HBE:16HBEge-G542X. We observed that CFTR function in each monolayer correlated with the percentage of wildtype cells. Importantly, previous studies have found that a population of 10% wildtype-expressing cells could achieve wildtype levels of function^15^. We found that integration rates of roughly 2% with the *CFTR* transgene-constructs achieved CFTR-specific inhibited currents at levels between the wildtype:G542X monolayers of 10% and 20% wildtype 16HBEs (**Figure 4H**).

## Discussion

Broad categories of nucleic acid-based gene therapies including antisense oligonucleotide (ASO), tRNA, mRNA, lentiviral vectors, and DNA-based therapeutics are of great interest for the treatment of CF and are currently under rapid advancement^37, 38^. However, there are limitations associated with these approaches, including but not limited (1) such approaches often only target a single mutations class, (2) lentiviral vector, though able to carry large cassettes are unable to target ABSCs as VSVG targets LDL receptors on the basolateral, but not apical side, (3) AAVs have limited packaging capacities and repeated dosing is largely inefficient, (4) AAVs trigger immune responses, and (5) common delivery mechanisms must penetrate the thick mucus layer and more^39^. Despite these limitations, it is crucial that a continued effort is made as gene editing of long-lived stem cell niches within affected tissues holds promise for sustained and robust treatment for CF patients. Recently, exciting data on platforms that leverage base editors or CRISPR-based correction of small mutations using single-stranded oligodeoxynucleotide (ssODN) donors have been published.^40, 41^ However, these approaches require the design and validation of reagents customized to address single mutations.

With this understanding, the primary goal of this work was to develop a nonviral gene-editing approach that could functionally and universally correct all CF-causing mutations and could be adapted for nanoparticle-based delivery. A candidate DNA HDR construct was designed to maximize the editing and functional efficacy of the constructs with an aim of overcoming nanoparticle-based delivery bottlenecks and ensuring straightforward encapsulation^42-46^. Ultimately, an AmC6 chemically modified ldsDNA with 50bp HA proved to be the most effective HDR donor. This donor design could be utilized more broadly for other diseases characterized by mutations in large genes exceeding the packaging limitation of AAV or research that requires a nonviral HDR donor for targeted site-specific integration.

The decreased integration efficiency of the full-length *CFTR* cDNA constructs demonstrate that further optimization of the strategy is necessary to achieve a higher proportion of allele integration events in the cell line population. Further optimization of these editing reagents to increase efficiency will be important when moving towards the less HDR prone primary ABSCs. This obstacle could partially be overcome by using LNPs rather than electroporation to deliver the editing reagents. LNPs have been demonstrated to have decreased cytotoxicity and induce fewer global changes to the transcriptome^47^.

We have minimized the size of the donor constructs to enable straightforward integration with nanoparticle-based delivery systems; however the size of these constructs still exceeds the size of DNA-based cargoes that have previously been delivered using LNPs. To bridge this gap, our team has parallel work recently conducted in which a LNP platform was optimized with the capacity to package and deliver these large ldsDNA constructs to the 16HBE cell line. Editing efficiencies modestly improved, but more encouragingly, expression and function equivalent to wildtype controls was achieved^48^.

However, delivery to primary ABSCs remains a major hurdle to *in vivo* treatment^49^. LNP administration via aerosolization or intravenous injection requires that nanocarriers traverse either the thick mucus layer, epithelial barrier, or endothelial barrier^50, 51^. Similar *ex vivo* strategies, namely autologous transplantation of edited ABSCs remains an unsolved challenge despite encouraging recent results^52^. Still, several exciting developments towards optimizing editing and delivery reagents are proving essential towards eventual clinical translation. Fortunately, with the reagents described and their compatibility with LNP-based delivery, this modular system can be tuned as necessary to address these challenges without altering the gene editing strategy.

Airway cell populations containing just 2% edited alleles restored approximately 50% of the functional activity of wildtype cells, which supports the field’s observations that low editing rates can yield therapeutically meaningful correction of the CF phenotype^21, 53^. As mentioned, codon-optimization techniques and BGH and WPRE sequences were included for their well-known capacity to boost expression, highlighting the necessity of regulatory elements in transgene design^54-61^.

Together, these results mark an important step towards a universal gene editing strategy to functionally replace the mutant *CFTR* gene. This research lays the foundation for a single, off-the-shelf therapy for virtually all CF patients, potentially offering hope to patients with currently untreatable mutations.

## Materials and Methods

### Cell Culture

T84 cells (ATCC; cat: CCL-248 Manassas, VA) were cultured in F-12K Hams Medium (ATCC; cat: 30-2004), with 10% fetal bovine serum (Omega Scientific FB-02 Tarzana, CA), and 1% penicillin/streptomycin/glutamate (ThermoFisher Scientific: cat: 10378016, Grand Island, NY). Passages were carried out weekly by detaching cells using 0.25% trypsin in EDTA (ThermoFisher cat: 25200114).

16HBE14o-cells were acquired from Millipore Sigma. 16HBEge-G542x cells were obtained via a materials transfer agreement between the Regents of the University of California, Los Angeles and Dr. Hillary Valley at the Cystic Fibrosis Foundation (CFF). 16HBE cell lines were cultured in Eagle’s Minimum Essential Medium (ATCC; cat: 30-2003) with 10% fetal bovine serum (Omega Scientific; cat: FB-02), 1% penicillin/streptomycin/glutamate (ThermoFisher Scientific: cat: 10378016), herein referred to as E10. Cells were cultured in flasks or well plates that were pre-coated with fetal bovine serum albumin (Millipore Sigma; cat: A7979 Saint Louis, MO), bovine collagen solution (Advanced Biomatrix cat: 5005 Carlsbad, CA), and fibronectin from human plasma (ThermoFisher cat: 33016015). Passages were carried out weekly by detaching cells using 0.25% trypsin in EDTA, quenching the reaction with E10, and splitting between 1:2 and 1:10.

### Donor Template Design and Generation

Double stranded DNA donor synthesis and amplification: Double-stranded DNA (dsDNA) donor fragments were synthesized as gBlocks by Integrated DNA Technologies (Coralville, IA). These gBlocks were subsequently cloned into plasmids using the TOPO Zero Blunt Cloning Kit (Thermo Fisher Scientific; cat: K2800J10). The dsDNA donors were then amplified by PCR from these plasmids with Platinum SuperFi II DNA Polymerase using the manufacturer’s protocol (ThermoFisher; cat: 12369010). Amplification utilized oligonucleotide primers that were modified at the 5’ end with AmC6, also supplied by Integrated DNA Technologies. The PCR products were then purified using 1X SPRI paramagnetic bead-based purification (Ampure XP; Beckman Coulter; cat: A63881 San Jose, CA). Following the first round of purification, the DNA was eluted twice with 150μL 1/10 TE in water. The 300µL total product was digested with Dpn1 at 37°C for 1hr to remove residual plasmid DNA. This was followed by a second round of purification, again with 1X SPRI beads. The final product was eluted in 80μL of pure TE.

### Delivery of RNP and HDR Donor

T84 cells were electroporated when cultures were 70-80% confluent. The day prior to transfection, 16HBE cells were split 1:2. For experiments with the full-length *CFTR* cDNA donors, 16HBE cells were cultured in 10μM Y-27632 ROCK inhibitor (StemCell Technologies: cat: 72304 Vancouver, Canada) starting from this split for the remainder of the experiment. 100pmol of purified Cas9-NLS protein (MacroLab, UC Berkeley, CA) was added to 120pmol sgRNA (Synthego Corporation, Menlo Park, CA) and let sit at room temperature for 20mins to form a ribonucleoprotein (RNP) complex. Cells were electroporated with the Lonza 4D-Nucleofector System (program FI 115 for T84s and CM 137 for 16HBEs) in SF Cell Line solution at a density of 200,000 cells per 20μL reaction with RNP, electroporation enhancer (IDT; cat: 1075916), and the HDR donor. Following electroporation, cells were let rest for 10mins then added to recovery plate with E10 medium and, if specified, 0.5μM AZD7648 (Selleckchem; cat: S8843 Houston, TX) and 0.5, 1, 2, or 3μM ART558 (MedChem Express; cat: HY-141520 Monmouth Junction, NJ) at various concentrations highlighted in the appropriate Figures.

### MTT Assay

Immediately after electroporation, 100μL of the 500μL of cells was seeded in a 96 well plate. 24 hours later, 10μL of MTT reagent (Biotium; cat: 30006 Fremont, CA) was added and incubated for 4hrs at 37C. Then, the formazan product was solubilized by adding 200μL DMSO and quantified by spectrometry at wavelength 570nm. Background was measured at wavelength at 630nm and subtracted from the on-target readings. Relative cell viability was normalized to the non-electroporated controls.

### Integration Analysis

Genomic DNA was extracted from edited cells for integration site analysis using the Invitrogen PureLink Genomic DNA Kit (Thermo fisher Scientific; cat: K182002) and quantified using the NanoDrop system (Thermo Fisher Scientific; cat: ND-2000). DNA samples were then analyzed by droplet digital PCR to measure integration rates. Two sets of primers were duplexed, each with their own fluorescent probe (FAM/HEX, Supplemental Table 2). To measure integration rates of the mCitrine reporter cassette, one primer was complementary to a *CFTR* gene sequence upstream from the left homology arm of the donor. The second primer bound to the mCitrine reporter cassette to allow for specific measurement of the integrated mCitrine transgene distinct from the endogenous *CFTR* gene. To measure integration rates of the *CFTR* donor cassette, one primer was bound to the bovine growth hormone polyA signal sequence to enable specific measurement of the integrated *CFTR* donor. The second primer was complementary to a *CFTR* gene sequence downstream from the right homology arm of the donor cassette. The FAM-conjugated nucleotide probe with a quencher also bound to the minus-strand DNA near the second primer. A reference primer/probe set was also delivered to recognize the *SDC4* gene on chromosome 20. 1μL of DraI endonuclease (New England Biolabs; cat: R0129S Ipswich, MA) was added to the reaction mixture (an enzyme that does not disrupt the experimental or reference amplicons) to reduce background. Each sample was digested at 37°C for 1hr before droplet generation with the Bio-Rad QX200 Droplet Generator (Bio-Rad, cat: 186-4002 Irvine, CA). The prepared samples were then assayed via the QX200 Bio-Rad Droplet Reader on the “Absolute” measurement setting (Bio-Rad; Cat: 186-40).

### Cutting Efficiency

The Cas9 cutting efficiency was measured via Inference of CRISPR Edits (ICE). PCR on genomic DNA extracted from edited cell populations was used to generate a 728bp amplicon for the 5’UTR guides (primer sequences: “AAAGCCGCTAGAGCAAATTT,” “TGTTGGCTGAATTCAGTCAA”) and 700bp for the Intron 1 guides (primer sequences: “GTCTGACAATTCCAGGCGCT,” “TTATTGATGCCTAGAGGGCAGA”. The resulting amplicon was Sanger sequenced and analyzed with the Synthego ICE web tool (Synthego Corporation, Menlo Park, CA).

### GUIDE-Seq

Double-strand DNA breaks at off-target cut sites were identified using genome-wide, unbiased identification of DSBs enabled by sequencing (GUIDE-Seq) using the protocol described by Malinin *et al.*^22, 23^. 16HBE or T84 cells were electroporated with Cas9-sgRNA RNP targeting the *CFTR* locus followed by the addition of 3μM or 5μM GUIDE-seq oligo. Sites of oligo trapping were enriched by locus-specific PCR followed by a second round of PCR to add index and adaptor sequences necessary for Next-Generation Sequencing. The amplicons were normalized using densitometry and pooled. The final library was quantified using ddPCR and sequenced using Illumina MiSeq. Data was analyzed following the bioinformatics pipeline from Malinin *et al.*^22, 23^.

### Western Blot

For immunoblots, cells were lysed in RIPA Lysis and Extraction Buffer (ThermoFisher Scientific cat: 89901 Grand Island, NY) with added HALT protease inhibitor (ThermoFisher Scientific; cat: 87786 Grand Island, NY) at a 1× concentration following the manufacturer’s protocols. Lysate concentrations were determined using the Pierce BCA protein assay (ThermoFisher Scientific; cat: 23227 Grand Island, NY) following the manufacturer’s protocol. Samples were treated for sodium dodecyl sulfate–polyacrylamide gel electrophoresis (SDS-PAGE) with NuPAGE LDS Sample Buffer (ThermoFisher Scientific; cat: NP0007) and NuPAGE Sample Reducing Agent (ThermoFisher Scientific; cat: NP0009), each to a 1× concentration. Lysates were diluted to contain 50μg of total protein for immunoblot gel loading to keep the total amount of protein loaded per lane constant to allow for valid loading controls. Note, CFTR protein becomes insoluble if the sample is heated above 60°C and will not enter the stacking gel. As such, the samples were denatured at 37°C for 20mins prior to loading into the stacking gel. Wild-type cells 16HBE14o- were used as a control to indicate the relative expression levels of CFTR protein. CFTR levels were detected using Ab UNC-596 (from J. Riordan, UNC^62^) at 1:500 in 5% milk in TBST. Protein quantification was assessed through densitometry via the ImageJ software. CFTR protein levels were normalized to the actin protein levels after quantification.

### Ussing Chamber Assay

16HBEge-G542X cells were seeded on 12mm polyester SnapWell inserts (Corning; cat: 3801 Corning, NY) coated in collagen type IV from human placenta (Millipore Sigma; cat: C5533) at a density of 500,000 cells/well and cultured in E10 in basolateral and apical chambers for at least 7 days prior to Ussing assays. Assays were performed using an EasyMount Ussing Chamber (Physiologic Instruments) at 37^°^C in HEPES buffered solutions with an imposed chloride gradient across the epithelia. The basolateral solution contained (mM): 137 NaCl, 4 KCl, 1.8 CaCl2, 1 MgCl2, 10 HEPES and D-Glucose, adjusted to pH 7.4 with NaOH/HCl ([Cl−]total: 146.6 mM). The apical solution was matched to the basolateral except for (mM): 137 Na-gluconate replaced 137 NaCl ([Cl−]total: 9.6 mM).

Electrode tips (Physiologic instruments) were partially filled with 3% LB Agar (Thermo Fisher; cat:22700041) in 3M KCl and backfilled with 3M KCl.

Cells grown on SnapWell inserts were mounted in the chambers and allowed to equilibrate for 10 – 15mins before recording baseline short-circuit current (Isc) for 10mins. At 10min intervals, 10 µM amiloride, 10 µM forskolin, 1 µM VX-770, and 20 µM CFTR-Inh172 were added sequentially to the apical and basolateral side of the chamber. At 10mins after the initial dose of CFTR-Inh172, a second dose of 20µM CFTR-Inh172 was added to the basal and apical chambers to extinguish any remaining current, and, 5mins later, 100µM adenosine triphosphate (ATP) was added to the basolateral and apical chambers which served as a quality control for the activation of calcium-activated Cl currents. I_sc_ was recorded for an additional 10 min before ending the experiment.

## Data Availability Statement

All data is provided in the main text or supplemental information. Raw sequencing data from GUIDE-Seq can be made available upon request.

## Acknowledgements

We thank the Cystic Fibrosis Foundation (CFF) for creating and generously sharing the 16HBEge-G542X cell line and for the CFTR Antibodies Distribution Program from which we received the UNC-596P CFTR antibody. We also thank California Institute for Regenerative Medicine (CIRM) Grant DISCO0-14458 (BNG, DBK, SJJ), the CFF Grant JONAS20XX0 (SJJ, DBK, BNG), and a New Horizons Award from the Cystic Fibrosis Research Institute, award #: 20213480 (SJJ, BNG, DBK) for providing the funds for this project.

## Author Contributions

Conceptualization: VS, PGA, BNG, SJJ, DBK,

Data curation: VS, PGA, CJJ, LEL, RAF

Formal analysis: VS, PGA, CJJ, LEL, RAF, JDL, BI

Funding acquisition: BNG, SJJ, DBK

Investigation: VS, PGA, CJJ, LEL, RAF, JDL, BI, ECD, NF

Methodology: VS, PGA, LEL, CJJ

Project administration: SJJ, BGN, DBK

Resources: VS, PGA, CJJ, LEL

Software: RAF, JDL

Supervision: VS, PGA, CJJ, LEL

Validation: VS, PGA, CJJ, LEL

Visualization: VS, PGA, CJJ, LEL

Writing – original draft: VS, CJJ

Writing – review & editing: VS, CJJ, PGA, LEL, RAF, RAS, BI, BNG, SJJ, DBK

## References

1. Riordan, JR, Rommens, JM, Kerem, BS, Alon, N, Rozmahel, R, Grzelczak, Z, et al. (1989). Identification of the Cystic-Fibrosis Gene - Cloning and Characterization of Complementary-DNA. Science 245: 1066–1072.

2. Guggino, WB, and Banks-Schlegel, SP (2004). Macromolecular interactions and ion transport in cystic fibrosis. Am J Respir Crit Care Med 170: 815–820.

3. Gustafsson, JK, Ermund, A, Ambort, D, Johansson, ME, Nilsson, HE, Thorell, K, et al. (2012). Bicarbonate and functional CFTR channel are required for proper mucin secretion and link cystic fibrosis with its mucus phenotype. J Exp Med 209: 1263–1272.

4. Cantin, AM, Hartl, D, Konstan, MW, and Chmiel, JF (2015). Inflammation in cystic fibrosis lung disease: Pathogenesis and therapy. J Cyst Fibros 14: 419–430.

5. Middleton, PG, Mall, MA, Drevinek, P, Lands, LC, McKone, EF, Polineni, D, et al. (2019). Elexacaftor-Tezacaftor-Ivacaftor for Cystic Fibrosis with a Single Phe508del Allele. N Engl J Med 381: 1809–1819.

6. Alameeri, A, Yavuz, BC, Lucca, F, Bambir, I, Famulska, P, and R, WFC (2025). Cystic fibrosis year in review 2024. J Cyst Fibros 24: 218–223.

7. Hussain, K, Karhana, S, Garg, A, and Khan, MA (2025). Efficacy of Trikafta (ELX/TEZ/IVA) & Symdeko (TEZ/IVA) in Treating Cystic Fibrosis with F508del Allele: A Systematic Review and Meta-analysis. Thorac Res Pract.

8. Clancy, JP, Cotton, CU, Donaldson, SH, Solomon, GM, VanDevanter, DR, Boyle, MP, et al. (2019). CFTR modulator theratyping: Current status, gaps and future directions. J Cyst Fibros 18: 22–34.

9. Graham, C, and Hart, S (2021). CRISPR/Cas9 gene editing therapies for cystic fibrosis. Expert Opin Biol Ther 21: 767–780.

10. Grieger, JC, and Samulski, RJ (2005). Packaging capacity of adeno-associated virus serotypes: impact of larger genomes on infectivity and postentry steps. J Virol 79: 9933–9944.

11. Song, Y, Lou, HH, Boyer, JL, Limberis, MP, Vandenberghe, LH, Hackett, NR, et al. (2009). Functional Cystic Fibrosis Transmembrane Conductance Regulator Expression in Cystic Fibrosis Airway Epithelial Cells by AAV6.2-Mediated SegmentalTrans-Splicing. Human Gene Therapy 20: 267–281.

12. Rose, I, Greenwood, M, Biggart, M, Baumlin, N, Tarran, R, Hart, SL, et al. (2025). Adenine base editing of CFTR using receptor targeted nanoparticles restores function to G542X cystic fibrosis airway epithelial cells. Cell Mol Life Sci 82: 144.

13. Geurts, MH, de Poel, E, Amatngalim, GD, Oka, R, Meijers, FM, Kruisselbrink, E, et al. (2020). CRISPR-Based Adenine Editors Correct Nonsense Mutations in a Cystic Fibrosis Organoid Biobank. Cell Stem Cell 26: 503–510 e507.

14. Bulcaen, M, Kortleven, P, Liu, RB, Maule, G, Dreano, E, Kelly, M, et al. (2024). Prime editing functionally corrects cystic fibrosis-causing CFTR mutations in human organoids and airway epithelial cells. Cell Rep Med 5: 101544.

15. Dorfman, R (2011). Cystic Fibrosis Mutation Database. Cystic Fibrosis Genetic Analysis Consortium: http://www.genet.sickkids.on.ca/Home.html.

16. Ostedgaard, LS, Rokhlina, T, Karp, PH, Lashmit, P, Afione, S, Schmidt, M, et al. (2005). A shortened adeno-associated virus expression cassette for CFTR gene transfer to cystic fibrosis airway epithelia. Proc Natl Acad Sci U S A 102: 2952–2957.

17. Vaidyanathan, S, Baik, R, Chen, L, Bravo, DT, Suarez, CJ, Abazari, SM, et al. (2022). Targeted replacement of full-length CFTR in human airway stem cells by CRISPR-Cas9 for pan-mutation correction in the endogenous locus. Mol Ther 30: 223–237.

18. Vaidyanathan, S, Kerschner, JL, Paranjapye, A, Sinha, V, Lin, B, Bedrosian, TA, et al. (2024). Investigating adverse genomic and regulatory changes caused by replacement of the full-length CFTR cDNA using Cas9 and AAV. Mol Ther Nucleic Acids 35: 102134.

19. Liu, X, Wang, X, Wu, X, Zhan, S, Yang, Y, and Jiang, C (2025). Airway basal stem cell therapy for lung diseases: an emerging regenerative medicine strategy. Stem Cell Res Ther 16: 29.

20. Berical, A, Lee, RE, Randell, SH, and Hawkins, F (2019). Challenges Facing Airway Epithelial Cell-Based Therapy for Cystic Fibrosis. Front Pharmacol 10: 74.

21. Johnson, LG, Olsen, JC, Sarkadi, B, Moore, KL, Swanstrom, R, and Boucher, RC (1992). Efficiency of gene transfer for restoration of normal airway epithelial function in cystic fibrosis. Nat Genet 2: 21–25.

22. Tsai, SQ, Zheng, Z, Nguyen, NT, Liebers, M, Topkar, VV, Thapar, V, et al. (2015). GUIDE-seq enables genome-wide profiling of off-target cleavage by CRISPR-Cas nucleases. Nat Biotechnol 33: 187–197.

23. Malinin, NL, Lee, G, Lazzarotto, CR, Li, Y, Zheng, Z, Nguyen, NT, et al. (2021). Defining genome-wide CRISPR-Cas genome-editing nuclease activity with GUIDE-seq. Nat Protoc 16: 5592–5615.

24. Kumar, P, Nagarajan, A, and Uchil, PD (2018). Analysis of Cell Viability by the MTT Assay. Cold Spring Harb Protoc 2018.

25. Shy, BR, Vykunta, VS, Ha, A, Talbot, A, Roth, TL, Nguyen, DN, et al. (2023). High-yield genome engineering in primary cells using a hybrid ssDNA repair template and small-molecule cocktails. Nat Biotechnol 41: 521–531.

26. Li, K, Wang, G, Andersen, T, Zhou, P, and Pu, WT (2014). Optimization of genome engineering approaches with the CRISPR/Cas9 system. PLoS One 9: e105779.

27. Yu, Y, Guo, Y, Tian, Q, Lan, Y, Yeh, H, Zhang, M, et al. (2020). An efficient gene knock-in strategy using 5’-modified double-stranded DNA donors with short homology arms. Nat Chem Biol 16: 387–390.

28. Selvaraj, S, Feist, WN, Viel, S, Vaidyanathan, S, Dudek, AM, Gastou, M, et al. (2024). High-efficiency transgene integration by homology-directed repair in human primary cells using DNA-PKcs inhibition. Nat Biotechnol 42: 731–744.

29. Dacquay, LC, Antoniou, P, Mentani, A, Selfjord, N, Martensson, H, Hsieh, PP, et al. (2025). Dual inhibition of DNA-PK and Polϴ boosts precision of diverse prime editing systems. Nat Commun 16: 4290.

30. Ranaghan, MJ, Li, JJ, Laprise, DM, and Garvie, CW (2021). Assessing optimal: inequalities in codon optimization algorithms. BMC Biol 19: 36.

31. Paremskaia, AI, Kogan, AA, Murashkina, A, Naumova, DA, Satish, A, Abramov, IS, et al. (2024). Codon-optimization in gene therapy: promises, prospects and challenges. Front Bioeng Biotechnol 12: 1371596.

32. Mauro, VP, and Chappell, SA (2014). A critical analysis of codon optimization in human therapeutics. Trends Mol Med 20: 604–613.

33. Grote, A, Hiller, K, Scheer, M, Munch, R, Nortemann, B, Hempel, DC, et al. (2005). JCat: a novel tool to adapt codon usage of a target gene to its potential expression host. Nucleic Acids Res 33: W526–531.

34. Raab, D, Graf, M, Notka, F, Schodl, T, and Wagner, R (2010). The GeneOptimizer Algorithm: using a sliding window approach to cope with the vast sequence space in multiparameter DNA sequence optimization. Syst Synth Biol 4: 215–225.

35. Valley, HC, Bukis, KM, Bell, A, Cheng, Y, Wong, E, Jordan, NJ, et al. (2019). Isogenic cell models of cystic fibrosis-causing variants in natively expressing pulmonary epithelial cells. J Cyst Fibros 18: 476–483.

36. Lesueur, LL, Mir, LM, and Andre, FM (2016). Overcoming the Specific Toxicity of Large Plasmids Electrotransfer in Primary Cells In Vitro. Mol Ther Nucleic Acids 5: e291.

37. Kreda, SM (2022). Oligonucleotide-based therapies for cystic fibrosis. Current Opinion in Pharmacology 66: 102271.

38. Allaire, NE, Griesenbach, U, Kerem, B, Lueck, JD, Stanleigh, N, and Oren, YS (2023). Gene, RNA, and ASO-based therapeutic approaches in Cystic Fibrosis. Journal of Cystic Fibrosis 22: S39–S44.

39. Lomunova, MA, and Gershovich, PM (2023). Gene Therapy for Cystic Fibrosis: Recent Advances and Future Prospects. Acta Naturae 15: 20–31.

40. Wei, T, Sun, Y, Cheng, Q, Chatterjee, S, Traylor, Z, Johnson, LT, et al. (2023). Lung SORT LNPs enable precise homology-directed repair mediated CRISPR/Cas genome correction in cystic fibrosis models. Nat Commun 14: 7322.

41. Sun, Y, Chatterjee, S, Lian, X, Traylor, Z, Sattiraju, SR, Xiao, Y, et al. (2024). In vivo editing of lung stem cells for durable gene correction in mice. Science 384: 1196–1202.

42. Kim, J, Vaughan, HJ, Zamboni, CG, Sunshine, JC, and Green, JJ (2021). High-throughput evaluation of polymeric nanoparticles for tissue-targeted gene expression using barcoded plasmid DNA. J Control Release 337: 105–116.

43. Eweje, F, Ibrahim, V, Shajii, A, Walsh, ML, Ahmad, K, Alrefai, A, et al. (2025). Self-assembling protein nanoparticles for cytosolic delivery of nucleic acids and proteins. Nat Biotechnol.

44. Patel, MN, Tiwari, S, Wang, Y, O’Neill, S, Wu, J, Omo-Lamai, S, et al. (2025). Safer non-viral DNA delivery using lipid nanoparticles loaded with endogenous anti-inflammatory lipids. Nat Biotechnol.

45. Zhu, Y, Shen, R, Vuong, I, Reynolds, RA, Shears, MJ, Yao, ZC, et al. (2022). Multi-step screening of DNA/lipid nanoparticles and co-delivery with siRNA to enhance and prolong gene expression. Nat Commun 13: 4282.

46. Liao, HC, Shen, KY, Yang, CH, Chiu, FF, Chiang, CY, Chai, KM, et al. (2024). Lipid nanoparticle-encapsulated DNA vaccine robustly induce superior immune responses to the mRNA vaccine in Syrian hamsters. Mol Ther Methods Clin Dev 32: 101169.

47. Vavassori, V, Ferrari, S, Beretta, S, Asperti, C, Albano, L, Annoni, A, et al. (2023). Lipid nanoparticles allow efficient and harmless ex vivo gene editing of human hematopoietic cells. Blood 142: 812–826.

48. Foley, R, Ayoub, PG, Sinha, V, Juett, C, Sonoyca, A, Duggan, EC, Lathrop, LE, Bhatt, P, Coote, K, Illek, B, Gomperts, BN, Kohn, DB, Jonas, SJ (2025). Lipid Nanoparticles for the Delivery of CRISPR/Cas9 Machinery to Enable Site-Specific Integration of CFTR and Mutation-Agnostic Disease Rescue. bioRxiv.

49. Wei, T, Sun, Y, Cheng, Q, Chatterjee, S, Traylor, Z, Johnson, LT, et al. (2023). Lung SORT LNPs enable precise homology-directed repair mediated CRISPR/Cas genome correction in cystic fibrosis models. Nat Commun 14: 7322.

50. Tafech, B, Rokhforouz, MR, Leung, J, Sung, MM, Lin, PJ, Sin, DD, et al. (2024). Exploring Mechanisms of Lipid Nanoparticle-Mucus Interactions in Healthy and Cystic Fibrosis Conditions. Adv Healthc Mater 13: e2304525.

51. Boucher, RC (2019). Muco-Obstructive Lung Diseases. N Engl J Med 380: 1941–1953.

52. Dorrello, NV, Guenthart, BA, O’Neill, JD, Kim, J, Cunningham, K, Chen, YW, et al. (2017). Functional vascularized lung grafts for lung bioengineering. Sci Adv 3: e1700521.

53. McCague, AF, Raraigh, KS, Pellicore, MJ, Davis-Marcisak, EF, Evans, TA, Han, ST, et al. (2019). Correlating Cystic Fibrosis Transmembrane Conductance Regulator Function with Clinical Features to Inform Precision Treatment of Cystic Fibrosis. Am J Respir Crit Care Med 199: 1116–1126.

54. Real, G, Monteiro, F, Burger, C, and Alves, PM (2011). Improvement of lentiviral transfer vectors using cis-acting regulatory elements for increased gene expression. Appl Microbiol Biotechnol 91: 1581–1591.

55. Donello, JE, Loeb, JE, and Hope, TJ (1998). Woodchuck hepatitis virus contains a tripartite posttranscriptional regulatory element. J Virol 72: 5085–5092.

56. Loeb, JE, Cordier, WS, Harris, ME, Weitzman, MD, and Hope, TJ (1999). Enhanced expression of transgenes from adeno-associated virus vectors with the woodchuck hepatitis virus posttranscriptional regulatory element: implications for gene therapy. Hum Gene Ther 10: 2295–2305.

57. Zufferey, R, Donello, JE, Trono, D, and Hope, TJ (1999). Woodchuck hepatitis virus posttranscriptional regulatory element enhances expression of transgenes delivered by retroviral vectors. J Virol 73: 2886–2892.

58. Levitt, N, Briggs, D, Gil, A, and Proudfoot, NJ (1989). Definition of an efficient synthetic poly(A) site. Genes Dev 3: 1019–1025.

59. Yew, NS, Wysokenski, DM, Wang, KX, Ziegler, RJ, Marshall, J, McNeilly, D, et al. (1997). Optimization of plasmid vectors for high-level expression in lung epithelial cells. Hum Gene Ther 8: 575–584.

60. Choi, JH, Yu, NK, Baek, GC, Bakes, J, Seo, D, Nam, HJ, et al. (2014). Optimization of AAV expression cassettes to improve packaging capacity and transgene expression in neurons. Mol Brain 7: 17.

61. Powell, SK, Rivera-Soto, R, and Gray, SJ (2015). Viral expression cassette elements to enhance transgene target specificity and expression in gene therapy. Discov Med 19: 49–57.

62. Cui, L, Aleksandrov, L, Chang, XB, Hou, YX, He, L, Hegedus, T, et al. (2007). Domain interdependence in the biosynthetic assembly of CFTR. J Mol Biol 365: 981–994.

