## Supplemental Materials for "Investigation of Double-Stranded DNA Donors and CRISPR-Cas9 RNP for Universal Correction of Mutations Causing Cystic Fibrosis in Human Airway Cells"

### Supplementary Materials

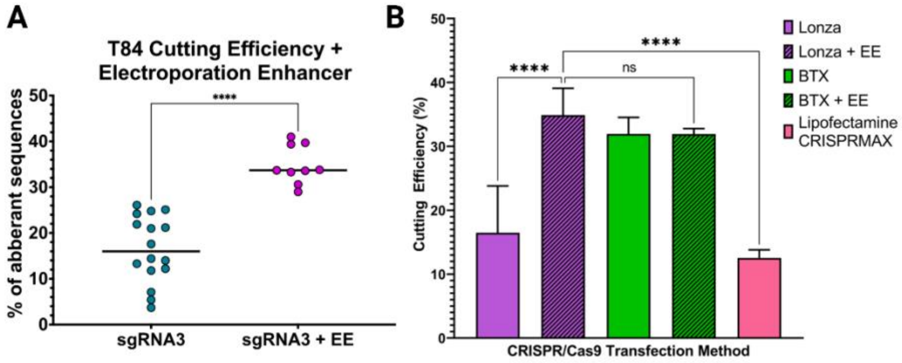


**Figure S1. Electroporation Enhancer Improves Delivery of sgRNA-Cas9 RNP**

(A) Cutting efficiency, measured by %indels, in T84 cells with or without electroporation enhancer (n=6). Each dot represents a different electroporated replicate (n=3). (B) Cutting efficiency following delivery through different platforms. Data is presented as means ± standard deviation. Statistical significance was analyzed using a one-way ANOVA followed by multiple paired comparisons for normally distributed data (Tukey test). All statistical tests were two-tailed and a p value of $< 0.05$, $^{**}p < 0.01$, $^{***}p < 0.001$, $^{****}p < 0.0001$.

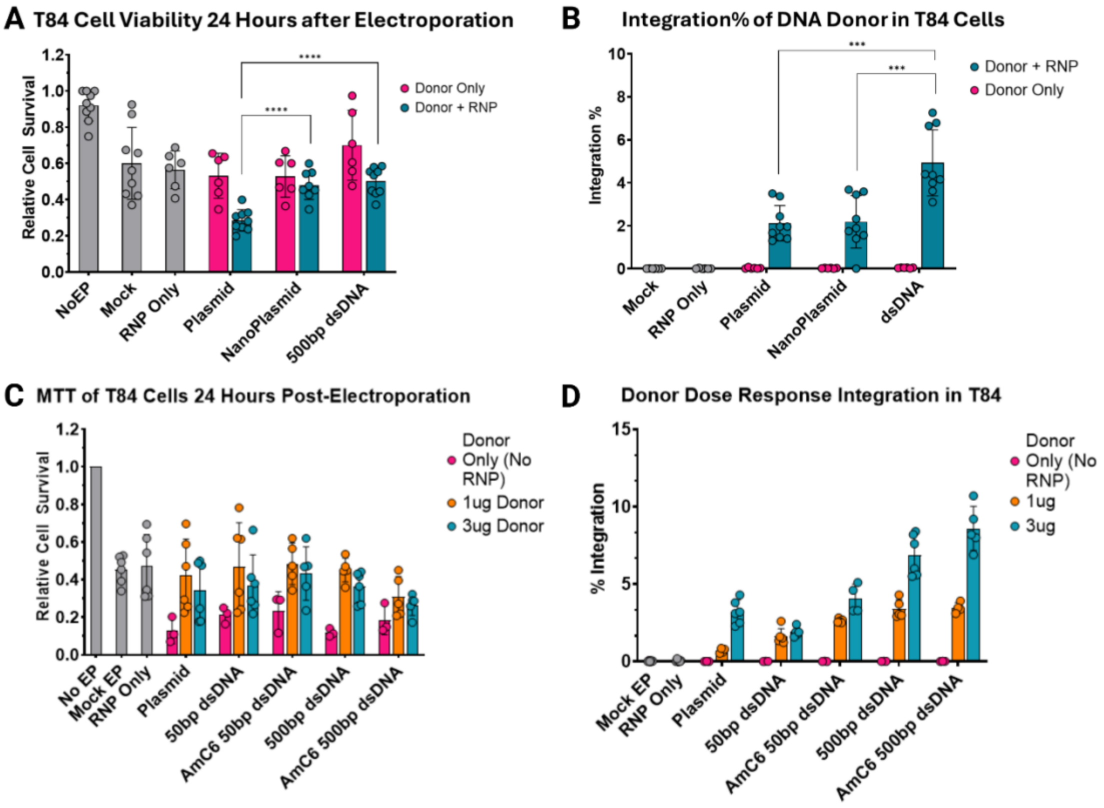


**Figure S2. Optimization of HDR Donors in the T84 Cell Line**

(A) Relative cell survival and (B) integration after electroporation delivering plasmid, nanoplasmid, or linear dsDNA HDR donor with 500bp HA construct to T84 cells (n=3). (C) Relative cell survival and (D) integration after electroporation delivering 50bp homology arm donors with or without AmC6 and 500bp donors with or without AmC6 to T84 cells (n=3). Statistical significance was analyzed using a one-way ANOVA followed by multiple paired comparisons for normally distributed data (Tukey test). All statistical tests were two-tailed and a p value of < 0.05, **p < 0.01, ***p < 0.001, ****p < 0.0001.

**Supplemental Table 1**

| sgRNA Name | Sequence | Location |
|---|---|---|
| sgRNA 3 | GCATGGTCTCTCGGGCGCTG | 5'UTR |
| sgRNA 13 | GCTAGTATATGATTATTTGG | Intron 1 |
| sgRNA 14 | CTTTGTCAAAGGGATTGGGA | Intron 1 |
| sgRNA 15 | AGTCAATTTCTATAAATACC | Intron 1 |
| sgRNA 17 | AAGACACGTGCCCACGAAAG | Intron 1 |
| sgRNA 18 | ACACGTGCCCACGAAAGAGG | Intron 1 |
| sgRNA 19 | CACGTGCCCACGAAAGAGGA | Intron 1 |

**Supplemental Table 2**

| Donor | Transgene Primers and Probe | Reference Gene Primers and Probe |
|---|---|---|
| 50bp HA mCitrine | F: AAGGAAGGGGTGGTGTG<br>R: cgaccaggatgggcac<br>P: TCTCTGACCTGCTGTGATGTC | F: AATACGCTGAGTGTCTCTTC<br>R: CTCACTCTCTTCAACGGG<br>P: TGAGAGGGCAGGGTCTGGGA |
| 500bp HA mCitrine | F: GGAGGGTCTAGGAAGCT<br>R: tcgcccttgctcacCAT<br>P: TCTCTGACCTGCTGTGATGTC | F: CTCTTCCAAAATACGCTGAG<br>R: GCAGAAGAATGACTCAATGC<br>P: TCCTCACTCTCTTCAACGGGT |
| 50bp HA CFTR cDNA | F: CAGCCATCTGTTGTTTGCC<br>R: CCAACCCATACACACGCC | F: AATACGCTGAGTGTCTCTTC<br>R: CTCACTCTCTTCAACGGG |

| | | |
|---|---|---|
| | P:<br>AGAGCCCACCGCATCCCCAG<br>(FAM) | P:<br>CCCACCGAACCCAAGAAACTAGAGGAGAAT<br>(HEX) |
